# MetaboXcan: A multiomic framework linking genetically predicted metabolites, gene expression, and complex traits

**DOI:** 10.64898/2026.07.15.738631

**Authors:** Festus Nyasimi, Sarah Sumner, Yanyu Liang, Xianyong Yin, Yoson Park, Andrew Brown, Sulaiman A. Lawal, Ana Viñuela, Lilian Fernandes Silva, Chunyu Liu, Sarah D. Slack, Erika Esquinca, Randi K. Johnson, Katerina Kechris, Markku Laakso, Michael Boehnke, Eric Fauman, Abraham A. Palmer, Sandra Sanchez-Roige, Ani Manichaikul, Nicholette D. Palmer, Hae Kyung Im

**Affiliations:** Section of Genetic Medicine, University of Chicago, Chicago, Illinois, United States of America; Department of Biostatistics and Center for Statistical Genetics, University of Michigan School of Public Health, Ann Arbor, Michigan, United States of America; Eli Lilly and Company, Indiana, United States of America; Department of Population Health and Genomics, School of Medicine, University of Dundee, Dundee, United Kingdom; Biosciences Institute, Faculty of Medical Sciences, University of Newcastle, Newcastle upon Tyne NE1 4EP, United Kingdom; Institute of Clinical Medicine, Internal Medicine, University of Eastern Finland, Kuopio, Finland; Department of Biostatistics, Boston University School of Public Health Boston, Massachusetts, United States of America; Department of Biomedical Informatics, University of Colorado Anschutz Medical Campus, Aurora, Colorado, United States of America; Department of Biostatistics and Informatics, University of Colorado Anschutz Medical Campus, Aurora, Colorado, United States of America; Department of Epidemiology, Colorado School of Public Health, Aurora, Colorado, United States of America; Internal Medicine Research Unit, Pfizer Worldwide Research, Development and Medical, Cambridge, Massachusetts, United States of America; Department of Psychiatry, University of California San Diego, La Jolla, California, United States of America; Institute for Genomics Medicine, University of California San Diego, La Jolla, California, United States of America; Department of Medicine, Division of Genetic Medicine, Vanderbilt University, Nashville, Tennessee, United States of America; Department of Genome Sciences, University of Virginia, Charlottesville, Virginia, United States of America; Department of Biochemistry, Wake Forest School of Medicine, Winston-Salem, North Carolina, United States of America

## Abstract

Genetically regulated molecular phenotypes, such as gene expression and metabolites, are widely thought to mediate the effect of disease-relevant genetic loci identified by genome-wide association studies (GWAS); however, molecular mechanisms connecting genetic variation to disease remain poorly understood. While transcriptome-wide association studies (TWAS) have identified disease-associated genes, strategies that integrate the metabolome remain underexplored. Here, we introduce MetaboXcan, a framework for predicting plasma metabolite levels from genetic data and associating them with complex traits using GWAS summary statistics. We trained lasso regression models on plasma metabolite and genotype data from the Metabolic Syndrome in Men Study (METSIM); these models outperform existing genetic metabolite predictors and generalize well across independent cohorts. MetaboXcan leverages these models to perform four complementary association analyses: (i) gene-to-trait (TWAS), (ii) gene-to-metabolite (M-TWAS), (iii) metabolite-to-trait (MWAS), and (iv) gene-expression–based metabolite-to-trait associations (g-MWAS). The framework further organizes results into metabolic pathways and gene–metabolite interaction networks to facilitate biological interpretation. Applied to chronic kidney disease (CKD), MetaboXcan identified known disease risk genes (*PDILT/UMOD, SPATA5L1/GATM*), as well as multiple CKD-relevant metabolites (e.g., glycine, homoarginine). Our integrated multiomic analysis enables biological interpretation by revealing glycine availability–centered biochemical axes spanning oxidative stress, cellular energetics, and vascular signaling, while nominating candidate mechanisms for downstream investigation. These results validate known disease-related biology with genetic evidence and generate new hypotheses for further investigation. MetaboXcan is publicly available and broadly applicable to any GWAS phenotype.

## Introduction

Understanding the genetic basis of complex traits and diseases is crucial for improving treatments and prevention strategies. While genome-wide association studies (GWAS) have identified thousands of genetic loci associated with human complex traits and diseases,^1^ translating these results into actionable discoveries remains challenging because most associated variants are in non-coding regions,^2^ obscuring their mechanistic function. Intermediate molecular phenotypes, such as gene expression and metabolites, may mediate the genetic effects on disease, motivating efforts focused on mapping genetic variation to these regulatory layers.

Significant progress has been made in identifying both gene- and metabolite-level mediators of genetic signals. For example, transcriptome-wide association studies (TWAS) link genetically predicted gene expression to complex traits, enabling the discovery of disease-relevant genes.^3,4^ Similarly, metabolite genome-wide association studies (mGWAS) have demonstrated that blood metabolites are heritable and influenced by genetic loci with shared disease and metabolic associations.^5^ Furthermore, thousands of metabolic- and disease-risk loci colocalize, implicating genes involved in metabolic regulation,^6–8^ suggesting putative etiological pathways for disease reflecting shared genetic architecture between genes and metabolites.

However, identifying loci jointly associated with both metabolites and disease requires large cohorts with matched metabolic and phenotypic data, which are costly and limited in availability.^9^ To overcome this limitation, we propose a TWAS-analogous approach—metabolome-wide association studies (MWAS)—which leverages genetic data to predict individual metabolomic profiles at scale and test for associations with disease using GWAS summary statistics. This approach enables population-level metabolic studies, improves statistical power by reducing the multiple testing burden, and extends applicability to contexts where direct observation of omics is impractical (e.g., brain regions, developmental stages). Genetic prediction also mitigates environmental noise inherent in omic datasets—critical for diseases where factors such as diet or chronic drug exposure can dramatically affect omics.

The MWAS approach requires a robust set of genetic prediction models for metabolites. While existing resources, such as OmicsPred,^10^ provide genetic predictors for a subset of metabolites, they lack an integrated multiomic framework for jointly interpreting metabolites, genes, and disease associations. Here, we investigate the genetic architecture of metabolites and train over 500 metabolite prediction models using matched genotype and plasma metabolite data from the Metabolic Syndrome in Men Study (METSIM)^8,11^. We show that these models improve prediction accuracy over existing approaches and generalize across independent cohorts and introduce MetaboXcan, a framework that integrates genetically predicted gene expression and metabolite levels in a suite of complementary association analyses. The framework further organizes associations within disease-associated metabolic pathways and integrated gene–metabolite networks, enabling the systematic interpretation of GWAS loci across regulatory layers and prioritization of candidate metabolites, genes, and pathways for downstream mechanistic studies.

To illustrate its utility, we applied MetaboXcan to chronic kidney disease (CKD), a well-studied complex trait characterized by the progressive impairment of kidney function. Although CKD GWAS have uncovered hundreds of associated loci, identifying causal genes and the molecular mechanisms linking them to disease remains challenging,^12–15^ making CKD an ideal test case for multiomic integrative analysis. Applied to CKD, MetaboXcan provides genetically grounded support for leading hypotheses in CKD pathogenesis and offers a structured framework for interpreting disease-associated genetic signals through integrated pathway and network analysis, enabling coherent multiomic hypothesis generation.

## Results

### MetaboXcan method overview

MetaboXcan is a framework that leverages multiple association analyses between gene expression, metabolites, and the trait of interest to nominate significant network clusters and elucidate associated metabolic pathways (Figure 1a). The model training component (Figure 1b) trains metabolite predictors from individual-level, observed plasma metabolite and genotype data and generates a metabolite prediction model and corresponding covariance matrix. The association analysi**s** component (Figure 1c) uses the trained molecular (i.e., gene and metabolite) predictors to perform four different analyses: (i) predicted gene expression-to-trait (**TWAS**), (ii) predicted gene expression-to-observed plasma metabolites (**M-TWAS**), (iii) direct predicted metabolite-to-trait (**MWAS**), and (iv) predicted metabolite-to-trait using aggregated predicted gene expression-to-trait associations (**g-MWAS**). The multiomic integration component (Figure 1d) comprises a metabolic pathway analysis and gene–metabolite network cluster analysis. The pathway analysis summarizes metabolite–trait associations (MWAS) by aggregating related metabolites into known metabolic pathways, highlighting the putative biochemical processes driving disease, while the network analysis maps trait-associated genes and metabolites into larger communities, illuminating possible shared biological processes.

**Figure 1:**
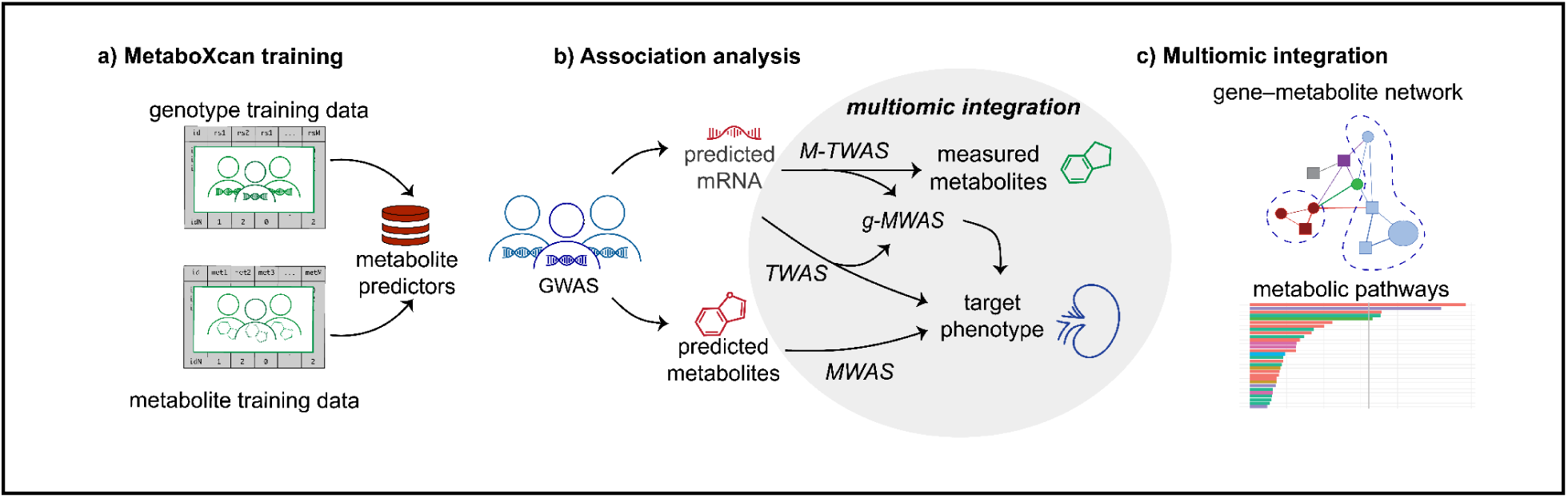
Overview of the MetaboXcan approach. a) MetaboXcan training. METSIM cohort genotype and measured metabolites are used to train models for specific metabolites. **c) Association analysis.** Application of MetaboXcan to a GWAS phenotype produces TWAS, g-MWAS, and MWAS association results. M-TWAS are agnostic to phenotype and provided with the software. The gene-to-trait association (TWAS) is performed with multi-tissue S-PrediXcan, i.e., the MultiXcan method. The metabolite-to-trait (MWAS) and gene-to-metabolite (M-TWAS) associations are performed using the same software as S-PrediXcan and MultiXcan, respectively. Gene expression–based metabolite-to-trait associations (g-MWAS) are calculated by aggregating TWAS p values for genes associated with each metabolite (see Figure 3). **c) Multiomic Integration.** Pathway analysis aggregates metabolites into chemically and functionally related groups. Gene–metabolite network clusters visualize relationships between species. Together these tools facilitate the biological interpretation of association results.

MetaboXcan is publicly available (https://github.com/hakyimlab/metaboxcan) and designed for broad phenotype applicability. A typical user will provide GWAS summary statistics, trait heritability estimated by Linkage Disequilibrium Score Regression (LDSC), and GWAS sample size. The software leverages precomputed metabolite and gene expression weights obtained from training (www.predictdb.org), along with metabolite pathway annotations (Metabolon) and an omics interaction network. MetaboXcan outputs gene–trait, gene–metabolite, and metabolite–trait associations, as well as metabolic pathways and gene–metabolite network clusters associated with the trait to aid biological interpretability.

### Metabolite training dataset

We used 6,136 participants with both genotype and metabolite data from Metabolic Syndrome in Men Study (METSIM)^8^ to train genetic predictors of metabolite levels, holding out 400 individuals for testing.

After quality control, the training dataset comprised 1,391 plasma metabolites profiled using untargeted relative quantitative liquid chromatography–tandem mass spectrometry (Metabolon DiscoveryHD4 platform). Untargeted metabolomics data often contain missing values, mostly due to metabolite concentrations below the limit of detection and random technical variations such as run day/time. In our dataset, a median of 234 metabolites were missing across samples (interquartile range [IQR] 196–332) and a median of 266 samples were missing across metabolites (IQR 11–1,894). We therefore imputed missing metabolite data using softImpute,^16^ which leverages correlations between metabolites.

### Metabolite levels are heritable and genetically predictable

To establish the feasibility of predicting metabolite levels from individual genotypes, we assessed the heritability of plasma metabolite levels using genome-wide complex trait analysis (GCTA).^17^ The heritability of the imputed metabolites ranged from 0–91% (median 17%; IQR 12–24%), with 91% (1,267/1,391) of the metabolites significantly heritable (*p* < 0.05; Figure 2a; Table S1). Imputed metabolites yielded slightly better heritability than metabolites with no imputation (median h^2^ = 0.17 and 0.16, respectively; Figure S1). In conjunction with many reported metabolite quantitative trait loci (mQTLs),^8^ these results suggest that many plasma metabolite levels are partially determined by genetic variation and are therefore likely to be genetically predictable.

**Figure 2.**
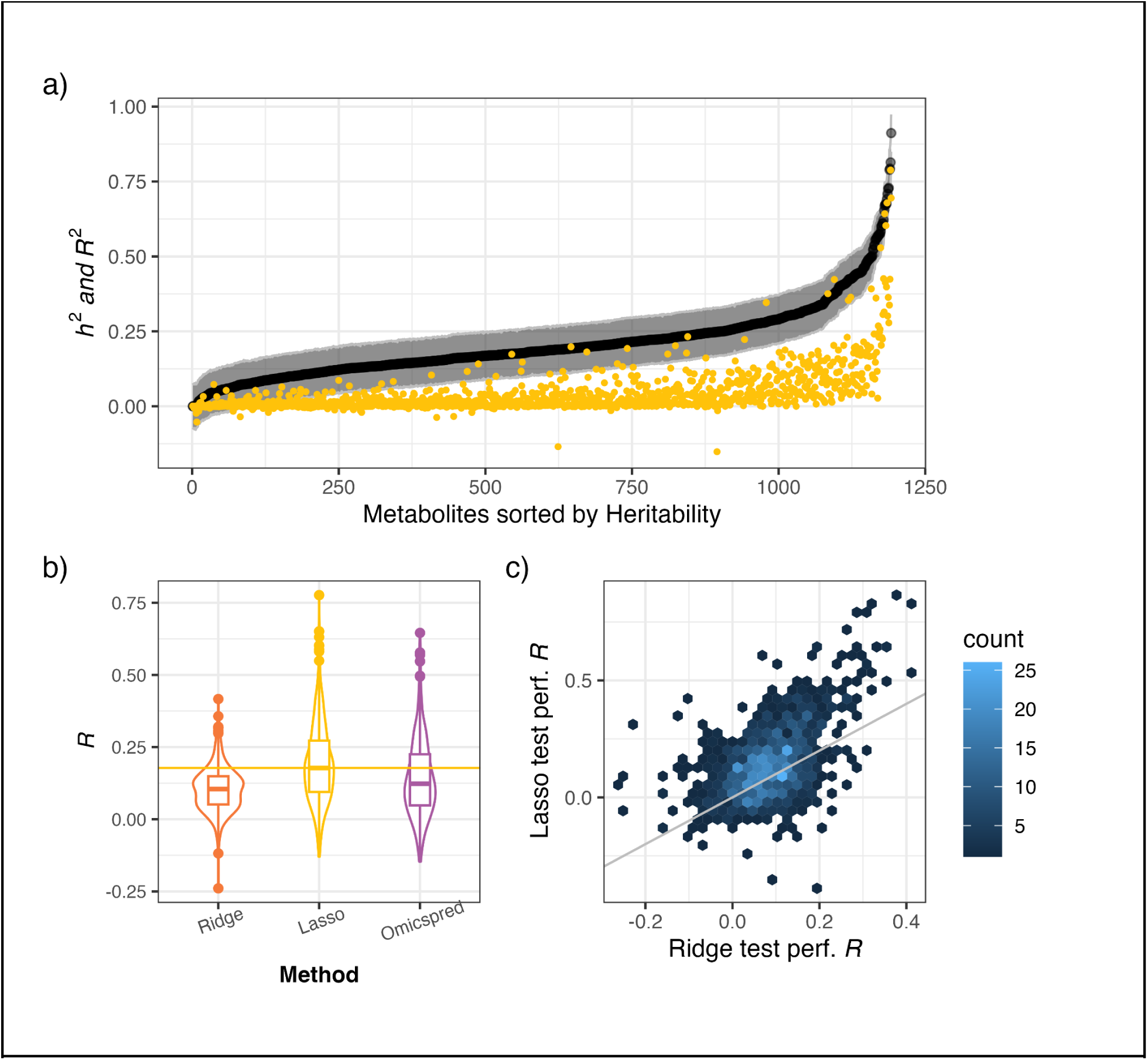
Heritability and predictability of metabolite levels. **a)** Heritability (h^2^, black) of observed plasma metabolites and Spearman correlation (R^2^, yellow) for observed and predicted metabolites using lasso prediction models in the METSIM held-out test set. The sign of the Spearman correlation coefficient was retained for R^2^. **b)** Box plot of Spearman correlation (R) between observed and predicted in the held-out test set for ridge and lasso models trained in METSIM and for OmicsPred models downloaded from omicspred.org. The yellow horizontal line indicates the median of the correlation for lasso, our model of choice. **c)** Comparison of lasso and ridge model performance (Spearman correlation, R) in a held-out test set. Both models were trained in the main METSIM data. Note that in panels b and c, only metabolites predicted by all compared models (lasso, ridge, and, in the case of b, Omicspred) are included for fair comparison.

To explore the underlying genetic architecture of metabolites, we trained linear predictors using both sparse and polygenic methods. We implemented genome-wide lasso regression (sparse) and genome-wide ridge regression (fully polygenic)^18^ on a single-nucleotide polymorphism (SNP) set restricted to Hapmap3.^19^ This restriction allowed us to keep the computational burden manageable while still capturing common variations across diverse human populations.

The lasso regression model generated predictors for 42% (580/1391) of metabolites with a Spearman correlation (R) between predicted and observed metabolite levels greater than 0.10 and p < 0.05 in the cross-validation set. In contrast, only 22% (301/1391) of the ridge regression predictors met these standards. Lasso regression yielded better predictors with a median R = 0.221 (min −0.390; IQR 0.162–0.321; max 0.888) in the held-out METSIM test set, whereas the ridge regression had a median R = 0.142 (min −0.158; IQR 0.087– 0.185; max 0.417) in the held-out test set. Both Spearman and Pearson correlation yielded similar results; here, we report Spearman correlation.

We compared our results to publicly available genetic predictors of metabolite levels (OmicsPred)^10^ developed using Bayesian ridge regression—another prediction method useful for highly polygenic traits. On average, the lasso predictors outperformed OmicsPred predictors for the held-out METSIM data (Figure 2b), despite being trained on a smaller sample size (6,136 vs. 8,153 individuals). The better performance of lasso predictors suggests that the genetic architecture of plasma metabolite levels is rather sparse, with relatively few large-effect variants rather than many small-effect ones.

Because our training set comprised male individuals with similar ancestry, we assessed the robustness of our predictors across ancestry and sex. To evaluate cross-ancestry performance, we applied our lasso-derived genetic predictors models to a cohort of individuals of Mexican descent from the Insulin Resistance Atherosclerosis Study–Classic (IRASC; Methods).^20,21^ Our results showed similar performance in IRASC (N=184) and METSIM (N=400) test sets (R = 0.42; *p* < 2.2 x 10^−16^; Figure S2), with the lasso predictors outperforming ridge and Omicspred models in the IRASC data.

We also compared the performance of our lasso model across sex in two distinct cohorts: the Insulin Resistance Atherosclerosis Study–Family Study cohort (IRASFS)^22,23^ and Genetic Epidemiology of Chronic Pulmonary Obstructive Disease cohort (COPDGene; Methods)^24,25^. Our predictors in both cohorts performed similarly across sex, with highly correlated prediction performance in sex-stratified subgroups (IRASFS: 408 metabolites, R = 0.67, *p* < 2.2 x 10^−16^; COPDGene: 405 metabolites, R = 0.74, *p* < 2.2 x 10^−16^; Figure S3). Together, these results demonstrate our models generalize well across ancestry and sex despite an all-male Finnish training cohort.

Finally, we evaluated the impact of imputation on genetic predictors trained using the lasso model by comparing predictors trained on observed data (no imputation) with those trained using mean imputation or softImpute. Using softImpute increased the number of well-predicted metabolites relative to no imputation (580 vs 543 well-predicted metabolites; R > 0.10) and yielded slightly higher-quality predictors than mean imputation, as reflected by consistently improved predictive performance relative to no imputation (Figure S4).

Based on these results, we used the lasso models trained on the softImpute-processed METSIM data for the remainder of our analysis. The prediction weights generated from training are shared with users through the MetaboXcan software.

### MetaboXcan association analysis: genes, metabolites, and trait

For any given GWAS phenotype, MetaboXcan determines associations between the trait and predicted gene expression (TWAS), predicted metabolites directly (MWAS), and metabolites indirectly using aggregated predicted gene expression (g-MWAS). An additional association component determines associations between predicted gene expression and observed metabolites (M-TWAS) in a phenotype agnostic analysis.

In a recent paper, we demonstrated that TWAS and other similar studies associating mediating traits (e.g., genes, metabolites) and polygenic traits are prone to statistical inflation—even when mediating traits are sparse—an effect that increases linearly with sample size and heritability of the polygenic trait.^18^ Accordingly, we corrected all association results for inflation using the variance control approach (Methods).

#### Gene expression-to-trait (TWAS)

MetaboXcan computes gene expression-to-trait associations with the traditional TWAS approach (i.e., correlating genetically predicted expression levels with a trait) applied across multiple tissues for increased power (MultiXcan; Methods).^26,27^ End users of MetaboXcan can use other gene expression models in PrediXcan format for prediction in different tissues and contexts, if desired. MetaboXcan also uses existing association tools that accept summary-level data (S-PrediXcan), facilitating the use of GWAS results in metabolite investigations.^26^

#### Gene expression-to-metabolite (M-TWAS)

For gene expression-to-metabolite associations, MetaboXcan uses precomputed associations agnostic to the target phenotype, which are included in the software package; however, users may substitute predictions from their preferred expression model when implementing the MetaboXcan framework.

To compute these associations, we performed a metabolite TWAS (M-TWAS) analysis treating each plasma metabolite as a trait and using gene expression prediction models from GTEx. Specifically, we predicted gene expression levels using genotype data in METSIM (6,136 individuals) and prediction models for the 49 GTEx tissues. We correlated these predicted values with the pre-processed observed plasma metabolite levels, and combined the results across tissues using the ACAT method—which increases power by accounting for dependency among individual tests.^28^ Despite minimal inflation due to the relatively small sample sizes, we adjusted our M-TWAS results in every tissue for inflation.

We identified a total of 13,037 significant gene-to-metabolite associations using a Bonferroni significance threshold (*p* < 2.3 x 10^−6^; 0.05/21,625). The genetically predicted expression showed associations with multiple metabolites, revealing widespread pleiotropic effects: each gene was associated with a median of two metabolites (IQR 1–4; max: 149).

#### Metabolite-to-trait (MWAS)

MetaboXcan calculates the association between genetically predicted plasma metabolite levels and the trait directly—an analysis we refer to as MWAS—using the S-PrediXcan framework with correction for inflation.

#### Gene expression–based metabolite-to-trait (g-MWAS)

To further boost power to detect trait-associated metabolites, MetaboXcan employs an indirect approach we call g-MWAS, which leverages the metabolite-to-gene and gene-to-trait associations to determine the metabolite-to-trait association. For each metabolite–trait pair, MetaboXcan identifies the genes linked to the metabolite and aggregates their gene-to-trait association *p* values using the ACAT method. Despite the correlated structure across genes, the ACAT method provides a “meta-analyzed” *p* value that serves as a proxy for the metabolite-to-trait association (Figure 3).

**Figure 3:**
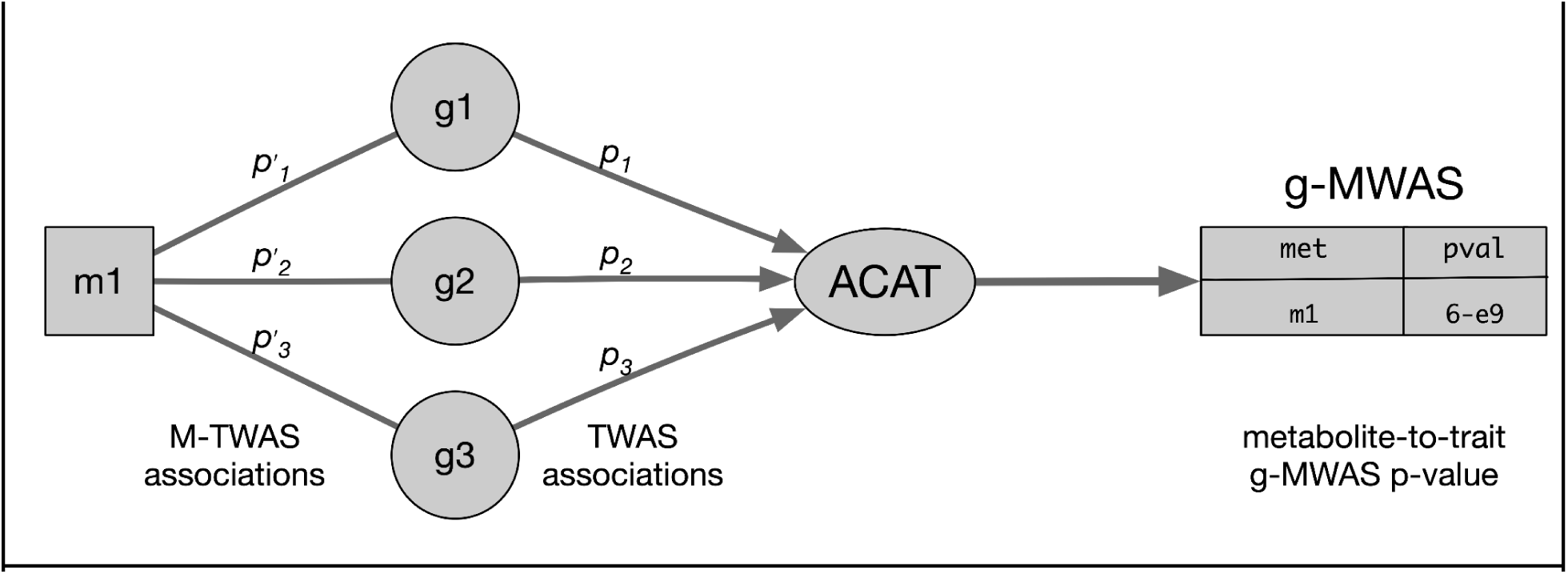
g-MWAS metabolite nomination workflow. Schematic representation of the gene expression–based metabolite-to-trait association (g-MWAS). For each metabolite, we identify the associated genes using M-TWAS associations and aggregate the TWAS p values of these genes using the ACAT method to obtain the g-MWAS p value.

### MetaboXcan multiomic integration: pathways and gene–metabolite network

Individual metabolite associations represent small components of a complex regulatory system. We hypothesized that by jointly analyzing genes and metabolites with pathway and network approaches (Figure 4), we could glean a more intuitive and comprehensive picture of the disease pathogenesis.

**Figure 4:**
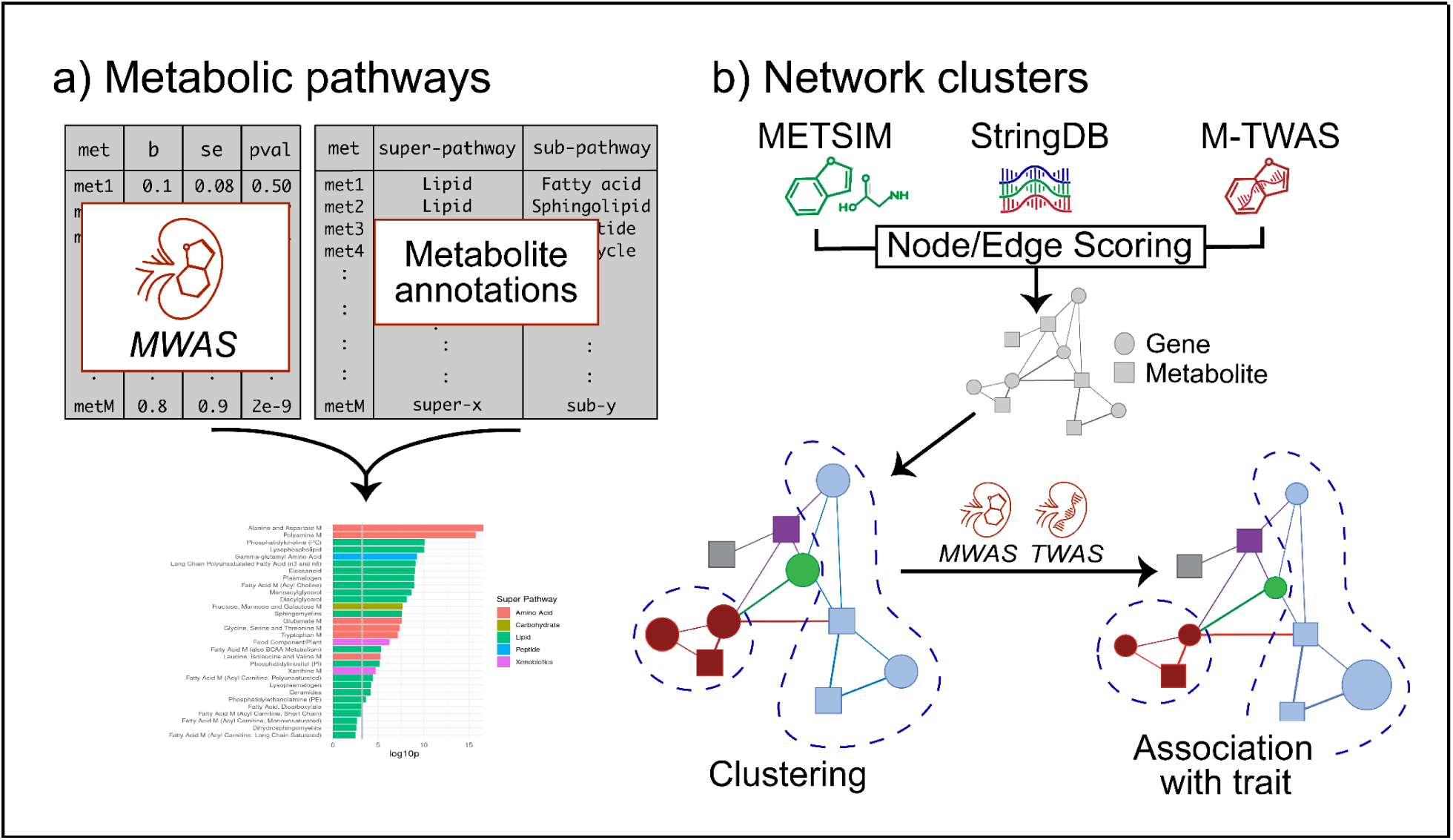
Network and pathway integration analysis. **a) Pathway analysis workflow.** MWAS association p values for metabolites annotated to the same pathway are aggregated via the ACAT method and pathways are tested for association with the target trait. **b) Network cluster workflow.** Genes are indicated as circles; metabolites as squares. Cluster membership is indicated by color and gene- or metabolite-to-trait association strength by shape size. The edges between metabolites are calculated using correlations between observed METSIM metabolite levels; edges between genes using STRING scores; and edges between genes and metabolites using predicted gene expression and metabolite associations (M-TWAS). Clusters are identified using the Louvain method and tested for association with the trait (Methods).

To implement the metabolite pathway analysis component, MetaboXcan aggregates metabolite-to-trait associations into biological pathways using annotations and categorizations provided by Metabolon HD4 and ranks them by calculating a combined *p* value for each pathway with the ACAT method.^28^ These pathway-level *p* values integrate evidence across all metabolites within a pathway, providing a single metric to assess the pathway’s relevance to the trait. Although Metabolon’s pathway annotations are typically based on chemical structure, users can infer functional assignments of each pathway by considering the shared biochemical roles of the pathway members.

The MetaboXcan network analysis generates an omics network consisting of gene–gene, gene–metabolite, and metabolite–metabolite interactions. Once MetaboXcan identifies the community clusters, it computes the significance of each cluster using the trait association *p* values of each node in the cluster with the ACAT method.

In designing the network (Methods), we used the STRING database’s functional protein–protein interactions to determine gene–gene interactions,^29^ Bonferroni-significant interactions from our M-TWAS analysis for the gene–metabolite interactions, and the METSIM cohort’s measured metabolite levels to calculate metabolite–metabolite associations. We chose the Louvain community detection approach to identify clusters because of its computational efficiency and high modularity.^30^ Other more statistically complex methods, such as Hotnet,^31^ require reducing the number of nodes to make them feasible, which we found too restrictive.

Integrating community cluster structure with the annotated pathway metabolites enables inference of coordinated biological processes that are obscured when considering individually significant genes and metabolites in isolation, providing a systems-level view of disease-relevant biology beyond what single-omics approaches can detect. To demonstrate this approach, we applied MetaboXcan to chronic kidney disease (CKD)—a complex trait with extensive GWAS and metabolomics literature that provides a rich framework for validation.

### MetaboXcan in chronic kidney disease: a model phenotype analysis

CKD is characterized by the progressive and long-term impairment of kidney function; although the pathogenesis of CKD is heterogeneous and incompletely characterized, converging evidence suggests several interacting factors including oxidative stress, mitochondrial dysfunction (cellular energetics), and endothelial dysfunction.^32–34^

We applied MetaboXcan to CKD using CKDGen GWAS summary statistics;^35^ full MetaboXcan association results for CKD are available in Table S2. Our results recapitulated known CKD-relevant genes and metabolites and revealed a biologically coherent pattern of associations in which the previously hypothesized pathogenic axes appear coordinated through glycine availability, as described below.

### TWAS implicates *PDILT/UMOD* and *SPATA5L1/GATM* loci in CKD

In CKD, the TWAS component identified four Bonferroni significant (p < 2.3 x 10^−6^, 0.05/21,360) genes: *PDILT* (p = 3.8 x 10^−9^), *SPATA5L1* (p = 2.3 x 10^−7^), and two unannotated long non-coding RNAs (lncRNAs): *RP11-96O20.5* (*p* = 3.8 x 10^−7^) and *EIF3J-AS1* (*p* = 5.7 x 10^−7^). Considering the locus-level ambiguity of TWAS associations that arises from linkage disequilibrium, the *PDILT* locus likely reflects the effect of *UMOD—*a kidney-specific gene flanking *PDILT* that is robustly associated with kidney function—although the precise mechanism linking this locus to CKD remains incompletely understood.^36^ Similarly, the *SPATA5L1* locus, including the two lncRNAs, likely captures the effect of *GATM,* which encodes arginine:glycine amidinotransferase (AGAT)—the rate-limiting enzyme in creatine synthesis and a source of homoarginine. *GATM* has been previously implicated in CKD.^37^

### MWAS and g-MWAS identify eleven significant metabolites in CKD

Direct metabolite association (MWAS) identified five Bonferroni-significant metabolites for CKD (Table 1): N-acetylglycine, glycine, *γ*-glutamylglycine, homoarginine, and propionylglycine. All of these metabolites have been previously implicated in CKD or related phenotypes.^38–41^ The complementary gene-expression–based analysis (g-MWAS) identified seven Bonferroni-significant metabolites, including homoarginine.

**Table 1.**
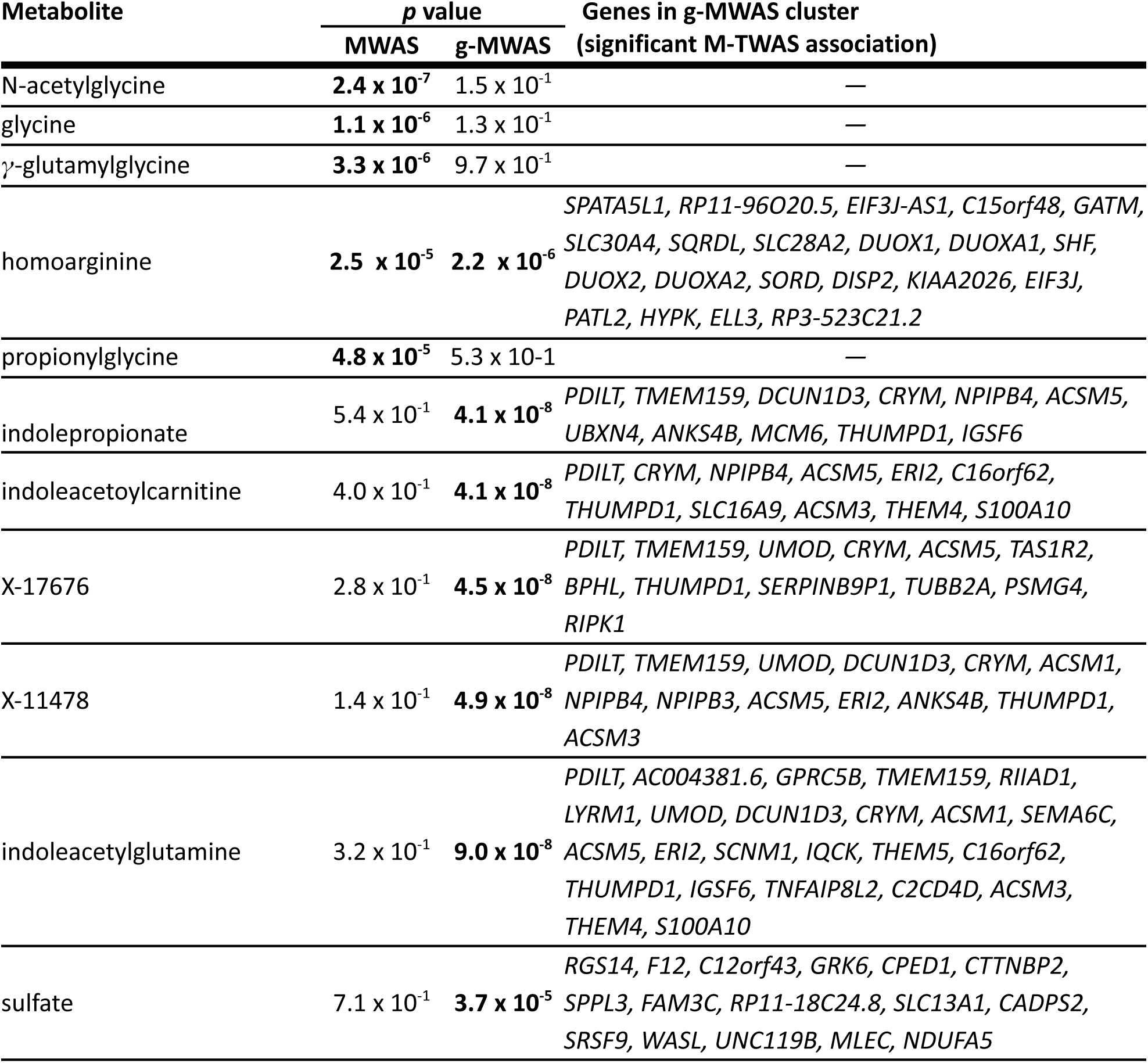
Significant MWAS and g-MWAS metabolites. Individual metabolites reaching Bonferroni significance (*p* < 8.6 x 10^−5^ 0.05/580; bolded *p* values) in MWAS and/or g-MWAS analyses. For g-MWAS significant metabolites, we list the genes with significant M-TWAS associations that contribute to the aggregated g-MWAS *p* value.

Examining the gene clusters driving the g-MWAS associations may give some insight into the CKD-related mechanisms involving these metabolites. For example, the gene clusters driving homoarginine and the top five g-MWAS metabolites (comprising three indoles and two uncharacterized metabolites) include *GATM-*locus and *PDILT/UMOD* genes, respectively. The homoarginine–*GATM* association is consistent with *GATM’*s established role in homoarginine biosynthesis and its previously reported association with CKD.^37^ Although *PDILT/UMOD* does not have an established connection with indole levels, gut-derived indole uremic toxins are cleared via proximal tubular organic anion transporters, suggesting that this association may reflect altered tubular function.^42^

Sulfate was Bonferroni significant in g-MWAS despite the absence of a Bonferroni-significant TWAS gene within its cluster. Instead, the signal was driven by two moderately associated genes, *RGS14* and *F12* (TWAS *p* values = 3.54 x 10^−6^ and 6.81 x 10^−6^, respectively), that share a locus with *SLC34A1*—a well-established kidney-function gene linked to proximal tubular integrity. Although the relationship between this locus and sulfate remains unclear,^15^ impaired renal sulfate reabsorption correlates with proximal tubular function,^43^ suggesting altered sulfate tubular clearance as a potential CKD mechanism.

Together, these findings demonstrate that MWAS and g-MWAS provide complementary and biologically interpretable signals, highlighting the utility of metabolite-level gene aggregation for pathway discovery and hypothesis generation.

### Pathway and network integration recapitulate glycine availability, oxidative stress response, cellular energetics, and vascular signaling in CKD pathogenesis

The pathway analysis revealed five significant pathways, which map to four biochemical roles consistent with known CKD pathogenesis: glycine availability, oxidative stress response, cellular energetics, and vascular signaling.

Although the Metabolon annotations provide a foundation for assessing the biochemical roles of metabolites, they do not always capture metabolite functions beyond the primary synthetic context. Therefore, we assessed the putative biochemical function of each pathway by considering both the Metabolon annotation and the canonical functions of the member metabolites found in relevant literature (Table 2). For example, of the metabolites assigned by Metabolon to glycine, serine, and threonine metabolism, all except N-acetylglycine are primarily involved in glycine availability through glycine biosynthesis and pool regulation (i.e., the control of free metabolite levels within the cell). Although annotated to glycine metabolism, N-acetylglycine is functionally a product of glycine conjugation with acetyl-CoA and aligns with other energy-related metabolites.^44^

**Table 2.**
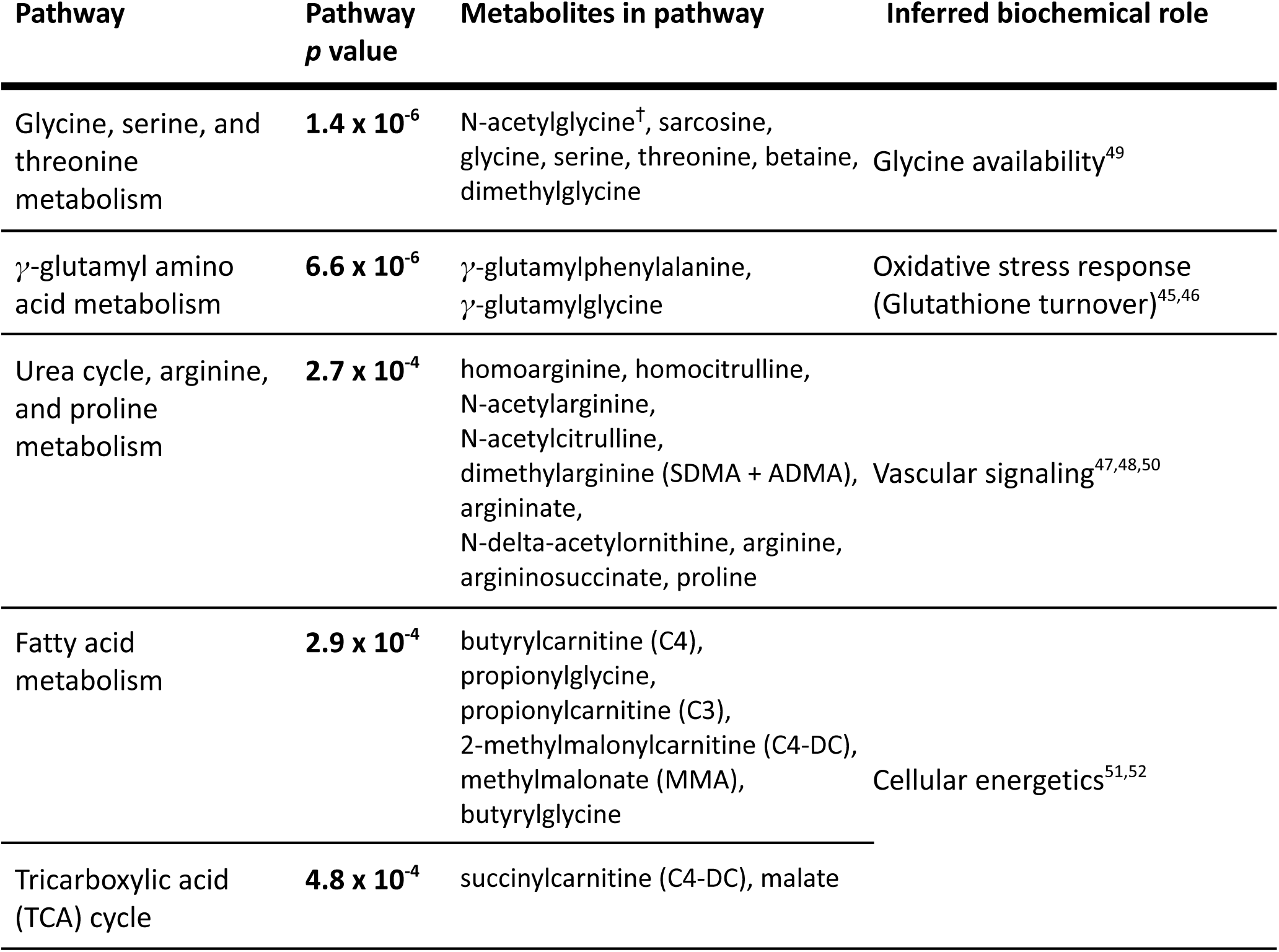
Significant pathways found with CKD pathway analysis. Pathways reaching Bonferroni significance (*p* < 6.0 x 10^−4^; 0.05/84; bolded *p* value). We list the metabolites annotated to the pathway and the biochemical role we inferred for the pathway. ^†^Based on literature, we interpreted N-acetylglycine as having a biochemical role in cellular energy, rather than glycine availability.

*γ*-Glutamyl amino acids are byproducts of the *γ*-glutamyl cycle, which is central to glutathione (GSH) homeostasis.^45^ GSH is a major scavenger of reactive oxygen species, which damage cellular macromolecules and contribute to renal fibrosis and CKD progression;^46^ the detection of *γ*-glutamyl amino acids suggests altered GSH turnover consistent with the known pathogenic role of oxidative stress in CKD.

The observed signal annotated as urea cycle, arginine, and proline metabolism pathway was primarily driven by homoarginine-related metabolites (homoarginine, homocitrulline, N-acetylcitrulline, and N-δ-acetylornithine), suggesting that the pathway-level association reflects perturbations in arginine-derived nitric oxide and endothelial signaling biology (vascular signaling) rather than canonical urea cycle flux. This is consistent with the prevailing view of homoarginine in CKD contexts.^47,48^

The metabolites annotated as fatty acid metabolism and TCA (tricarboxylic acid) cycle pathways all contribute cellular energy function. Biochemically, the *β*-oxidation of fatty acids and the TCA cycle are tightly linked pathways integral to energy production in the mitochondria.

In the network analysis for CKD, most individually significant metabolites and genes (i.e., MWAS, g-MWAS, and TWAS results) cluster closely, with glycine occupying a central position (Figure 5a). Despite the complexity of the full network, comprising ∼22,000 genes and metabolites organized into over 270 clusters (Supplemental data), the significant species are tightly clustered and highly interconnected, providing reassurance that we are capturing cohesive biologically meaningful interactions.

**Figure 5.**
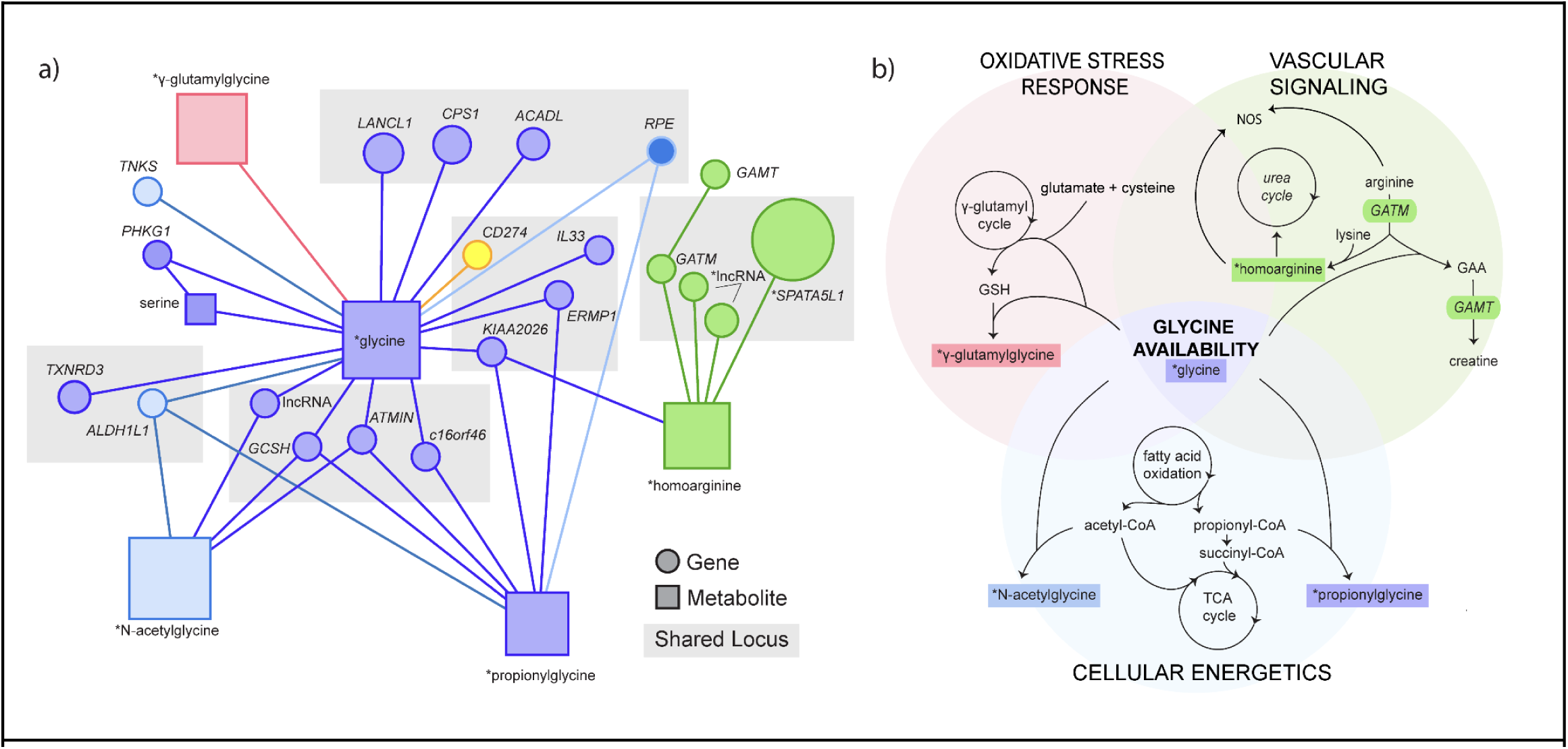
Network integration and pathway analysis results for CKD. **a) Portion of the network for CKD centered on glycine.** Inclusion criteria: i) all first-degree nodes for glycine, ii) all Bonferroni TWAS, MWAS, or g-MWAS significant species within three nodes of glycine, iii) shared locus with a TWAS-significant gene meeting criterion ii. Although GAMT does not meet these criteria, it is included to illustrate the relationship of GATM to creatine synthesis captured by the network. Genes are indicated as circles; metabolites as squares. The grey shading indicates genes at a shared locus. * indicates individually Bonferroni-significant genes and metabolites. Shape color indicates network membership, while shape size indicates the z-score. All clusters shown are at least nominally associated (p < 0.05) with CKD: Cluster 42 (blue, *p* = 2.1 x 10^−6^), Cluster 91 (purple, *p* = 2.8 x 10^−6^), Cluster 13 (red, *p* = 4.4 x 10^−4^), and Cluster 16 (green, *p* = 2.3 x 10^−3^). **b) Biochemical relationship of significant pathways and metabolites.** Schematic representation of canonical biochemical relationships between significant pathways and metabolites and genes in significant network clusters. Colors of genes (ovals) and metabolites (rectangles) match network cluster assignments in panel (a). Glutathione (GSH); guanidinoacetate (GAA).

All of the clusters containing MWAS-significant metabolites or TWAS-significant genes are also at least nominally significant in CKD. In fact, sulfate is the only individually significant metabolite or gene that does not reside in a significant cluster (Cluster 56, *p* = 1) or connect closely to another individually significant species.

Although not visualized in Figure 5a, the *PDILT/UMOD* and N-acetylglycine belong to the same cluster, separated by multiple intermediate nodes (Figure S5). The metabolites in the g-MWAS significant indole cluster (including the two uncharacterized metabolites) are all directly connected to *PDILT/UMOD.* The distant clustering with N-acetylglycine suggests that *PDILT/UMOD* may be broadly related to energy metabolism, but not a direct influence on N-acetylglycine levels.

To determine whether these signals reflect coherent biological processes, we mapped the roles of individually significant metabolites and genes appearing in the network clusters to the biochemical functions identified above (Figure 5b). This integrated analysis revealed that significant metabolites and genes cluster in patterns mirroring three canonical biochemical axes implicated in CKD: oxidative stress response, cellular energetics, and vascular signaling—with glycine availability emerging as a common upstream determinant.

Glycine is centrally located in the portion of the network containing CKD-significant genes and metabolites; it is directly connected to *γ*-glutamylglycine and shares at least one genetic locus each with N-acetylglycine, propionylglycine, and homoarginine. Notably, glycine is a required component in the synthesis of all the MWAS-significant metabolites—with the exception of homoarginine. However, homoarginine synthesis is indirectly dependent on glycine availability; the AGAT enzyme that produces homoarginine also synthesizes creatine in a parallel, glycine-dependent pathway. The dependence of multiple significant metabolites on glycine availability positions it at the center of the other biochemical processes identified by MetaboXcan, relationships reflected in the network clusters.

Glycine is also required for GSH synthesis. As previously discussed, *γ*-glutamylglycine is produced during GSH turnover and likely indicates oxidative stress response in CKD. The direct connection between glycine and *γ*-glutamylglycine in the network reinforces this relationship and suggests that glycine availability may influence the oxidative stress response axis mediated by GSH turnover.

N-acetylglycine and propionylglycine (both acylglycines) can form when acyl-CoAs (compounds produced by fatty acid oxidation and consumed by the TCA cycle) conjugate with glycine. This process serves as a buffering mechanism against the oxidative stress induced by excessive accumulation of acyl-CoAs when energetic flux is disrupted.

In our network, N-acetylglycine and propionylglycine connect to glycine through multiple mutually associated (M-TWAS) genetic loci, which may reflect a disruption in cellular energetic flux, the glycine-mediated acyl-CoA detoxification process, or both. Indeed, the functional roles of these genetic loci suggest an intersection between glycine availability, oxidative stress response, and cellular energetic axes. Genes in these loci have established functional roles in regulating glycine availability (*ALDH1L1* and *GCSH*)^53,54^ or immune-mediated oxidative stress response pathways (*ALDH1L1, TXNRD3,* and *ATMIN*)^55–57^.

In addition, glycine and propionylglycine both associate with a locus comprising *CPS1*, *ACADL, LANCL1* and *RPE*, which have established roles in mitochondrial metabolism—including nitrogen handling, fatty acid oxidation, and redox balance^58–62^. Importantly, *ACADL*^59^ and *LANCL1*^61^ are both highly expressed in the kidney and have been implicated in CKD pathogenesis.

Furthermore, glycine, propionylglycine, and homoarginine share an association with a locus containing *CD274 (PD-L1)*, *IL33,* and *ERMP1*—genes highly expressed in the kidney and with enriched immune and vascular signaling functions.^63,64^ As outlined in the pathway analysis, homoarginine plays a critical role in vascular function and endothelial health, while propionylglycine is a byproduct of dysregulated energetic axes. Taken together, this mutual association suggests vascular stress and energy axes are related to glycine availability in CKD pathogenesis and that these axes may be further mediated by immune response—a possibility that warrants further investigation.

Although not directly associated with glycine, the *SPATA5L1/GATM* locus associates strongly with homoarginine in our network. *GATM* is a well-known CKD risk locus thought to impact kidney function primarily through either creatine availability or vascular signaling^37,65,66^. While we cannot definitively rule out that the *GATM–*homoarginine signal reflects creatine availability (creatine is not predicted by MetaboXcan), homoarginine is consistently found as significantly associated with CKD across analyses (i.e., MWAS, g-MWAS, pathway, and network cluster), suggesting the *GATM*–homoarginine cluster reflects a vascular signaling axis rather than creatine-related energy pathways.

Overall, MetaboXcan results in CKD identify glycine as a central metabolic hub connecting three biochemical axes: oxidative stress, cellular energetics, and vascular signaling. These findings genetically validate leading hypotheses in CKD pathogenesis and demonstrate that MetaboXcan can uncover coherent biological architecture for complex traits—including signals not detectable by single-omics approaches—highlighting the value of integrated interpretation of genetically predicted multiomics in investigating disease pathogenesis.

## Discussion

While GWAS effectively identify loci associated with disease, they offer no information about the molecular pathways connecting genotype and phenotype. In addition to gene expression, metabolite levels can serve as intermediate molecular phenotypes and provide insight into the etiology of complex traits; however, widespread metabolic profiling is impractical. MetaboXcan predicts plasma metabolite and gene expression levels associated with complex traits using genetic predictors, providing an efficient and broadly applicable strategy for nominating molecular links between genetic variation and traits. The association module of MetaboXcan, based on the PrediXcan family of tools,^26^ can be used with individual- and summary-level data and incorporates multiple tissues for gene-level predictions. Due to the limited availability of individual-level data in most cases, we optimized the association module to use GWAS summary statistics. MetaboXcan’s analyses focus on identifying genes, metabolites, and metabolite pathways and clusters closely associated with the trait and potentially important to etiology. The interaction network clusters closely related genes and metabolites associated with the trait and the pathway analysis aggregates associated metabolites into candidate metabolic pathways, which not only increases statistical power to detect associations, but points to plausible causal mechanisms. This integrative approach, which combines evidence from different levels of molecular associations (genes, metabolites, and pathways), should deepen our understanding of the genetically driven molecular mechanisms underlying disease and facilitate the identification of biomarkers and potential therapeutic targets.

MetaboXcan improves on previous attempts to study imputed metabolite levels from genetic data. One study included a relatively small cohort of individuals (n=291) and metabolites (n=338).^67^ In contrast, our larger training cohort resulted in a greater number of higher-quality metabolite predictors: after filtering for prediction performance, we retained 580 well-predicted metabolites from thousands of individuals, a significant improvement over the previous study. Our trained metabolite models also retained performance with test data in a different ancestry and with balanced gender, despite using an all male training set with a homogeneous ancestral background. Additionally, our lasso models slightly outperformed the OmicsPred models in an independent test set, despite OmicsPred being trained on a larger cohort (8,153 European individuals vs. 6,136 in METSIM) and covering more metabolites (726 vs. 580), suggesting model choice can outweigh training set size in some circumstances.

A comparison between our M-TWAS results and a prior metabolite-level GWAS in the same cohort (METSIM) demonstrates the power and reliability of M-TWAS. The metabolite GWAS identified 2,030 significant single variant associations.^8^ Through fine-mapping and knowledge-based approaches, they nominated known genes for 1,375 of these independent variants. In contrast, our M-TWAS approach detected 13,037 gene–metabolite pairs, of which 703 overlapped with the 1,357 high-confidence pairs reported in the METSIM study. Because putative causal genes nominated by the fine-mapping and knowledge-based methods are considered high-confidence candidates, the recovery of these associations underscores the robustness of our approach. Further supporting this robustness, we observed extensive pleiotropy for genes associated with metabolites, especially those involved in lipid and amino acid pathways—a pattern consistent with the GWAS findings.

We applied MetaboXcan to CKD to evaluate the model’s ability to integrate genetically predicted gene regulation and metabolites into a biologically interpretable framework for identifying structured relationships between genes and metabolites in a disease context. In CKD, MetaboXcan’s individual association results for genes and metabolites (i.e., TWAS, MWAS, and g-MWAS) genetically validate associations found in the literature, but to our knowledge the identification of glycine availability as a central metabolic hub for multiple genetically mediated biochemical axes is a novel finding. Because the metabolite structure in the pathway enrichment and network analysis mirrors established biochemical relationships, we propose the genetically inferred associations capture coordinated metabolic programs rather than isolated signals. Importantly, the interdependent axes of oxidative stress response, cellular energetics, and vascular signaling align with established hypotheses for CKD,^32–34^ and multiple independent studies provide orthogonal support for the central role of glycine availability.^38–40,68^ Together, these results demonstrate the utility of an integrative framework for recovering biologically meaningful structure from genetic data and informing future mechanistic studies.

While MetaboXcan’s CKD results are promising, our method has some inherent limitations. First, our method does not establish causality between the complex trait and predicted metabolites or pathways. Although the method mitigates concerns regarding reverse causation and environmental confounding by leveraging genetically predicted metabolites and genes, additional lines of evidence are needed to infer causal relationships between metabolites, metabolic pathways, and traits.

The metabolomic training data were collected from Finnish men aged 45–74 years, which may limit the models’ ability to predict genetically influenced metabolic states that manifest at different life stages or under alternative environmental exposures. Further, because our study is restricted to the 580 (104 of which are uncharacterized) metabolites we could reliably predict using untargeted metabolomics, metabolites or pathways relevant to specific phenotypes may not be represented in our models due to analytical constraints or tissue specificity, and uncharacterized metabolites preclude assignment of implicated signals to known biochemical pathways.

As more robust datasets become available, we will update and improve the prediction models, which will enhance the reliability of the network analysis and identified candidate pathways. We also expect these improvements to further increase the cross-ancestry portability of the model and resolve the cohort-specific issues addressed above. In the meantime, the g-MWAS analysis helps address this power issue by providing an additional line of evidence: the proxy gene–trait associations provide useful information on pathways associated with the trait, even when the MWAS result is not significant, as evidenced by the indole and sulfate findings in CKD. Until more robust training sets are available, we can leverage this association to improve the power to detect trait-associated metabolites.

Finally, our pathway analysis relies on the accuracy of the available metabolite annotations. Many metabolites are relevant in multiple interconnected metabolic pathways (e.g., glycine), but the current annotation classifies them under a single super-pathway, and in some instances, the annotation files may be incomplete or inaccurate. Further, as seen in the case of N-acetylglycine in our CKD association analysis, Metabolon annotation is largely based on biosynthetic pathway or molecular class, and does not necessarily capture all relevant functions of each metabolite. We emphasize the importance of inspecting the metabolites that drive the pathway associations and considering the composition of the metabolite group when interpreting the pathway results.

Given the reliance of our method on inferred associations, independent replication and validation are essential for novel findings. *In silico* replication could be achieved by repeating MetaboXcan with metabolite prediction models trained with independent cohorts, which would assess the robustness and generalizability of the identified associations. Additional validation could be obtained through gene knockout studies and/or consistent associations with directly profiled metabolites from relevant tissues.

In conclusion, our study demonstrates the benefit of integrating different molecular intermediate phenotypes (i.e., genes and metabolites) to identify known and novel disease-relevant associations. Additionally, the high-quality genetic predictors for metabolites can be used to infer association with traits using existing methods. While our findings provide genetic support for metabolic pathways and molecular networks that may inform disease pathogenesis, they should be interpreted in light of the limitations of the study.

## Methods

### Materials availability

This study was performed in silico and did not generate any new reagents.

### Dataset utilised for training and testing

#### METSIM data used for training the models

The Metabolic Syndrome in Men Study (METSIM) was a cohort study of 10,197 Finnish men (aged 45–74 years at baseline) that investigated factors (including plasma metabolites) that are associated with type 2 diabetes and cardiovascular diseases.^8,11,69^ Array genotyping for all participants was performed with the Human OmniExpress BeadChip (OmniExpress) and Infinium HumanExome BeadChip (exome array) platforms. Sampling was done on participants who were neither diagnosed with diabetes nor were taking diabetes medications. Only 6,136 participants that passed various quality control filters^8^ were selected for metabolomic profiling. Non-targeted metabolomics profiling was performed on plasma samples at Metabolon which resulted in 1,391 detected and quantified metabolites. Out of the 1,391 detected metabolites 1,099 were characterized, named and categorized into super and sub-pathways while 292 remained uncharacterized. We used this dataset to train the genetic predictors of metabolites.

#### IRAS-FS data used for testing

The Insulin Resistance Atherosclerosis Family Study (IRAS-FS), a cohort of Mexican descent and with a balanced female-to-male ratio, was designed to investigate the genetics and epidemiology of glucose homeostasis.^22,23^ The IRAS-FS is one of the cohorts participating in Genetics Underlying Diabetes in Hispanics (GUARDIAN) Consortium^70^. Genotyping was done using the Illumina OmniExpress Array. Metabolite profiling was done on fasting plasma samples collected at baseline and stored at −80°C. Profiling was performed using untargeted liquid chromatography–mass spectrometry (DiscoveryHD4 panel) by Metabolon resulting in 1274 quantified metabolites.

We selected 1,027 individuals from this dataset whose genotype data and plasma metabolite measurements were both available. We were able to match 815 metabolites and used this data set as an independent test set for general model performance and performance across sexes.

#### IRASC data used for testing

The Insulin Resistance Atherosclerosis Study Classic (IRASC) dataset (n=184) was also one of the cohorts participating in the GUARDIAN Consortium.^20,21,70^ It has a similar ancestry profile as the IRAS-FS and was genotyped using the same platform. Metabolite profiling was performed using untargeted liquid chromatography–mass spectrometry by Metabolon. The IRASC provides individual-level data (genotype and matched plasma metabolites, n=815). We used IRASC as an independent test set to evaluate the performance of our trained models.

#### COPDGene data used for testing

We used the Genetic Epidemiology of Chronic Pulmonary Obstructive Disease (COPDGene) cohort^24,25^ as part of our performance evaluation of METSIM-trained models across sex. COPDGene represents never-, current-, and former-smoker adults at risk for developing COPD. Plasma metabolic profiling and genotyping was completed previously for a subset of COPDGene Phase 2 participants.^25^ We included 953 non-Hispanic white participants—487 male (51%), 748 former-smoker (78.5%), 205 current-smoker (21.5%)—with available metabolomic, genetic, and covariate data.

We adjusted the observed metabolite measures for age, sex, smoking status, pack years, BMI, and post-bronchodilator FEV1 percent predicted using a linear model. Genotyping was done with the HumanOmniExpress array, imputed to the TOPMed R3 panel after standard quality control, and SNPs with imputation R^2^ > 0.3 and MAF > 0.01 were input to the METSIM-trained prediction models. We used the Pearson correlation to compare the residuals of the adjusted observed metabolites to the genetically predicted metabolites for the 405 metabolites available in both.

#### Publicly available GWAS for application

We applied the MetaboXcan framework using publicly available GWAS summary statistics from CKD.^35^ We used a uniform harmonization, format standardization, missing data imputation, and other quality assurance steps as described in Barbeira et al.^26^

### Metabolite prediction model training

#### Training genotyping and quality control (METSIM)

We performed quality control using Plink software^71^ on the imputed individual-level genetic data. Individuals with missingness >1% were excluded, and variants departing from the Hardy–Weinberg equilibrium (HWE) (p < 1 x 10^−6^) or with missingness > 1% were removed. The genotypes were filtered to select only variants with minor allele frequency (MAF) of > 0.01. These SNPs were further filtered to select only non-ambiguous SNPs which are in the HapMap3 dataset (A list of variants was extracted from CEU individuals in HapMap3). The resulting set of SNPs were used for downstream computations.

We calculated genetic principal components from the quality controlled genotype using Plink and GCTA software.^17,71^ The genetic principal components were used as covariates.

#### Training metabolite measure processing and quality control (METSIM)

The 1,391 profiled metabolite levels from METSIM individuals were already processed whereby covariates (age at sampling, Metabolon batch, and, for lipid traits only, usage of lipid-lowering medication) were regressed out, and the residuals were inverse normalized.^8^ We imputed missing metabolites using the softImpute as described below.

#### Imputation of missing metabolites

The preprocessed metabolite levels contained some missing values. Missingness was mostly attributable to metabolite concentrations below the limit of detection and random technical variations. To resolve the missingness in the metabolite measures we used the softImpute package in R.^16^ The method leverages the underlying pattern and relationships within the available data to make informed estimates of the missing data.

As a first step we performed a grid search to find the best rank to use for imputation: we removed 20% of the data from the metabolite matrix—this included both missing data and available data. Using the new dataset, we ran an imputation and then calculated the MSE of each rank using the imputed matrix and the original matrix. The rank with the minimum MSE was used for final imputation of the metabolite matrix.

#### Heritability estimation

We estimated the heritability of each metabolite using linear mixed models and calculated variances using restricted maximum likelihood (REML) as implemented in GCTA. The model was set to only output h^2^ estimates between 0 and 1. We constructed the genetic relatedness matrix (GRM) based on total SNPs that passed the quality check filters and selection process. We also estimated heritability using Bayesian Sparse Linear Mixed Model (BSLMM)’s linear mixed model.^79^

#### Building prediction models from individual-level data

We built metabolite prediction models for each metabolite from individual-level data using two methods: ridge regression and lasso regression, both of which cover different assumptions. We split the training data into a training set (n=5736) and testing set (n=400). The testing set was only used for testing the final models generated.

We performed ridge regression with all the pre-selected genome-wide SNPs (HapMap3 set) for each metabolite; it used the L2 regularization parameter, and retained all the SNPs used. We used lasso regression to select optimal SNPs associated with each metabolite; it used the L1 regularization parameter to perform SNP selection. The lasso regression was implemented in the Snpnet R package.^72^ Lasso regression performs feature selection by imposing a penalty on the absolute value of the coefficients, resulting in sparse models. On the other hand, ridge regression shrinks the coefficients towards zero, resulting in simple models which are less likely to overfit the training data. We evaluated the models for their performance, and selected the optimum model.

We trained and tested each metabolite prediction model on the corresponding test dataset to determine the correlation and R^2^ between the actual and predicted metabolite levels. We used the calculated R^2^ and the h^2^ estimates to filter metabolite models to be included in the final best model.

#### Evaluating model performance

We evaluated the prediction performance of all models on the METSIM test set and the IRASC dataset. We used the prediction models to impute the metabolite level from the genotype data and then compared the imputed metabolite level and the observed metabolite levels. We evaluated the prediction performance using Spearman correlation R and R^2^. We also evaluated the fivefold, cross-validated performance of each model with the Spearman’s between the observed and predicted metabolite levels for each fold and averaged across folds.

### Molecular trait to phenotype associations

#### MetaboXcan association using summary statistics (TWAS and MWAS)

We used GWAS summary statistics to infer the association between genes (TWAS) and metabolites (MWAS) with the GWAS phenotype. This was implemented in MetaboXcan using S-PrediXcan,^26^ which uses the pretrained gene and metabolite prediction models, model covariance files, and the GWAS summary statistics.

#### Statistical adjustment for inflation

Liang et al.^18^ demonstrated that TWAS and related methods suffer from statistical inflation when the target complex trait is highly polygenic—even when the mediating traits (here, metabolites) are sparse. A simple example illustrates the source of the inflation: assume that the target trait (Y) is fully polygenic, meaning every variant has a nonzero effect on Y; also assume that an unrelated molecular trait has a single QTL. In this case, the TWAS association will capture an association between the target trait and that QTL—even in the absence of a causal relationship between the molecular and target traits—because for a fully polygenic target trait, the QTL has an actual direct effect on the trait.

Liang et al. quantified the inflation caused by this polygenicity-driven pleiotropy by deriving a mathematical expression for the Eχ^2^ statistic of the association in a more general context. They showed that the effect increases linearly with the sample size of the GWAS (N) and the polygenic heritability of the target trait (h^2^).

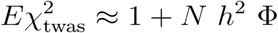

The Φ parameter is a complex function of the effect size of causal variants for the mediating trait that can be empirically estimated using actual genotype-based null polygenic trait simulations and predicted mediating traits. Heritability of the target trait can be estimated from the GWAS summary statistics using the Linkage Disequilibrium Score Regression (LDSC) method.^72^

Accordingly, we calibrated our MetaboXcan results using the newly developed inflation adjustment method. Using UK Biobank individual-level data, we estimated the metabolite-specific inflation parameter Φ and incorporated it into the model database. We calculated the sample size for each trait and estimated heritability using LDSC using the Hapmap 3 set of SNPs (as used for the estimation of the factor Φ). Once we gathered all necessary parameters (Φ, N, and h^2^), we calibrated our results using the variance parameter ΦNh^2^. We calibrated the z-score using the variance control approach as Z/sqrt(1 + ΦNh^2^).

### Gene to metabolite association (M-TWAS)

For gene-to-metabolite associations, we used the gene prediction models from GTEx to impute gene expression levels using the METSIM individual-level data. This was followed with association testing between the predicted gene expression and the measured metabolite levels across all 49 tissues using PrediXcan. We calibrated our results using the variance control method described above. This was followed by aggregating the tissue level results using the ACAT method. We only retained Bonferroni-significant associations, which we used downstream to define gene–metabolite edges in our base network.

### Omics network integration and cluster detection

We defined the gene and metabolite interaction networks using different approaches. For the development of gene–gene network we tested three approaches: using a GTEx co-expressing and co-regulation network alone, using the protein–protein interaction network downloaded from the STRING v11.5 database (i.e., functional protein association networks; https://string-db.org),^29^ and using an ensemble approach by combining the networks to find the best network. The protein–protein network was robust in our analysis. Because it provides additional information about the genes and contains confidence scores for each of the edges, we selected it for the gene–gene network. We normalized the protein scores and retained only those edges with scores exceeding the 99.5th percentile to develop our network. The protein–protein interaction network was then annotated with the specific gene names and information.

To develop the metabolite–metabolite interaction network, we used the measured metabolite levels from the METSIM cohort. We correlated all metabolite pairs and filtered out pairs with a correlation R < 0.2. The R values were used as edge weights in the network. For gene–metabolite edges we used the Bonferroni significant M-TWAS association and converted the *p* values into edge scores using a sigmoid function.

The gene–gene, metabolite–metabolite, and gene–metabolite interactions formed our omics network, which we used to perform cluster detection using the Louvain community detection algorithm. Once we detected community clusters, we selected all member nodes for each cluster and retrieved their trait association *p* values from either TWAS or MWAS association analysis. We then used the ACAT method to compute a combined *p* value for each cluster. Clusters were ranked based on these combined *p* values.

### Metabolite pathway analysis

The metabolites measured were annotated into super- and subpathways according to Metabolon HD4 annotations. We used the ACAT method^28^ to combine the *p* value of association for all the metabolites in a single pathway to find how strongly the pathway associated with a given trait. We ranked the pathways and interpreted Bonferroni significant pathways as associated with the trait of interest.

## Data and Code availability

The MetaboXcan software, the CKD analysis report and the CKD interactive network are available at https://github.com/hakyimlab/metaboxcan.

Metabolomics data is available at Metabolomics Workbench (https://www.metabolomicsworkbench.org/) with project identifier PR000907.

## Web Resources

Training and test-set data are available at the following locations.

METSIM, IRASC, and IRASFS datasets are available through the NIH dbGaP database: https://www.ncbi.nlm.nih.gov/gap/.

OmicsPred: https://www.omicspred.org/.

COPDGene genetic data is available through the COPDGene Data Coordinating Center. (https://copdgene.org; study specifics: https://copdgene.org/P3/COPDGeneMoP_Phase3_Version_14_2023_04_10.pdf).

## Acknowledgements

This research was conducted using the UK Biobank Resource under Application Number 89052. We utilized resources at the Argonne Leadership Computing Facility, a DOE Office of Science User Facility, supported under Contract DE-AC02-06CH11357. Additional computational support was provided by the University of Chicago’s Research Computing Center and Beagle3. We also acknowledge resources from the Center for Research Informatics, funded partially by the Biological Sciences Division at the University of Chicago. Finally, we thank Natalia Gonzales and Frederick Naumann for their editorial feedback.

## Author Contributions

Conceptualization: FN, HKI; Methodology: FN, YL, HKI; Software: FN, YL, HKI; Validation: FN, YP, CL, SDS, EE, RKJ, KK, EF, AM, HKI; Formal Analysis: FN, SS, YL, HKI; Data Curation: FN, SS, HKI; Writing – Original Draft: FN, SS, HKI; Writing – Review & Editing: FN, SS, YP, AV, EF, AP, SSR, AM, HKI; Visualization: FN, SS; Supervision: HKI; Project Administration: HKI; Resources: XY, AB, SAL, AV, LFS, CL, SDS, EE, RKJ, KK, ML, MB, NP

## Declaration of generative AI and AI-assisted technologies in the manuscript preparation process

During the preparation of this work some of the authors used Claude and ChatGPT in order to improve grammar and flow of the paragraphs. After using this tool/service, the authors reviewed and edited the content as needed and take full responsibility for the content of the published article.

## Supplemental Information

### Supplemental Figures

**Figure S1:**
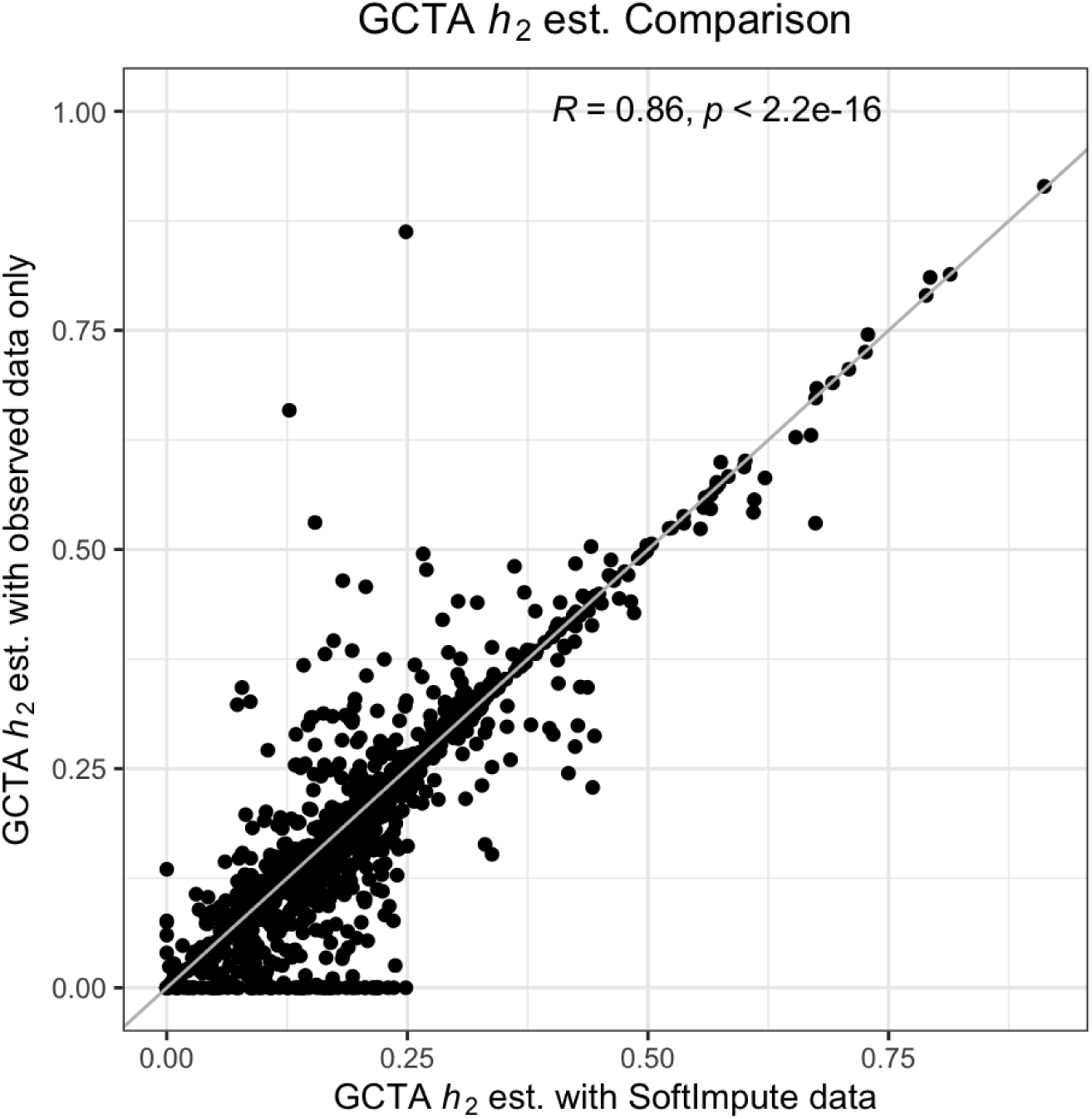
Heritability values of imputed (softImpute) and observed data. Comparison of heritability estimate (h^2^) of metabolites between observed-only and softImpute-imputed data. The softImpute yielded higher heritability estimates: softImpute median heritability = 0.17, observed median heritability = 0.16.

**Figure S2:**
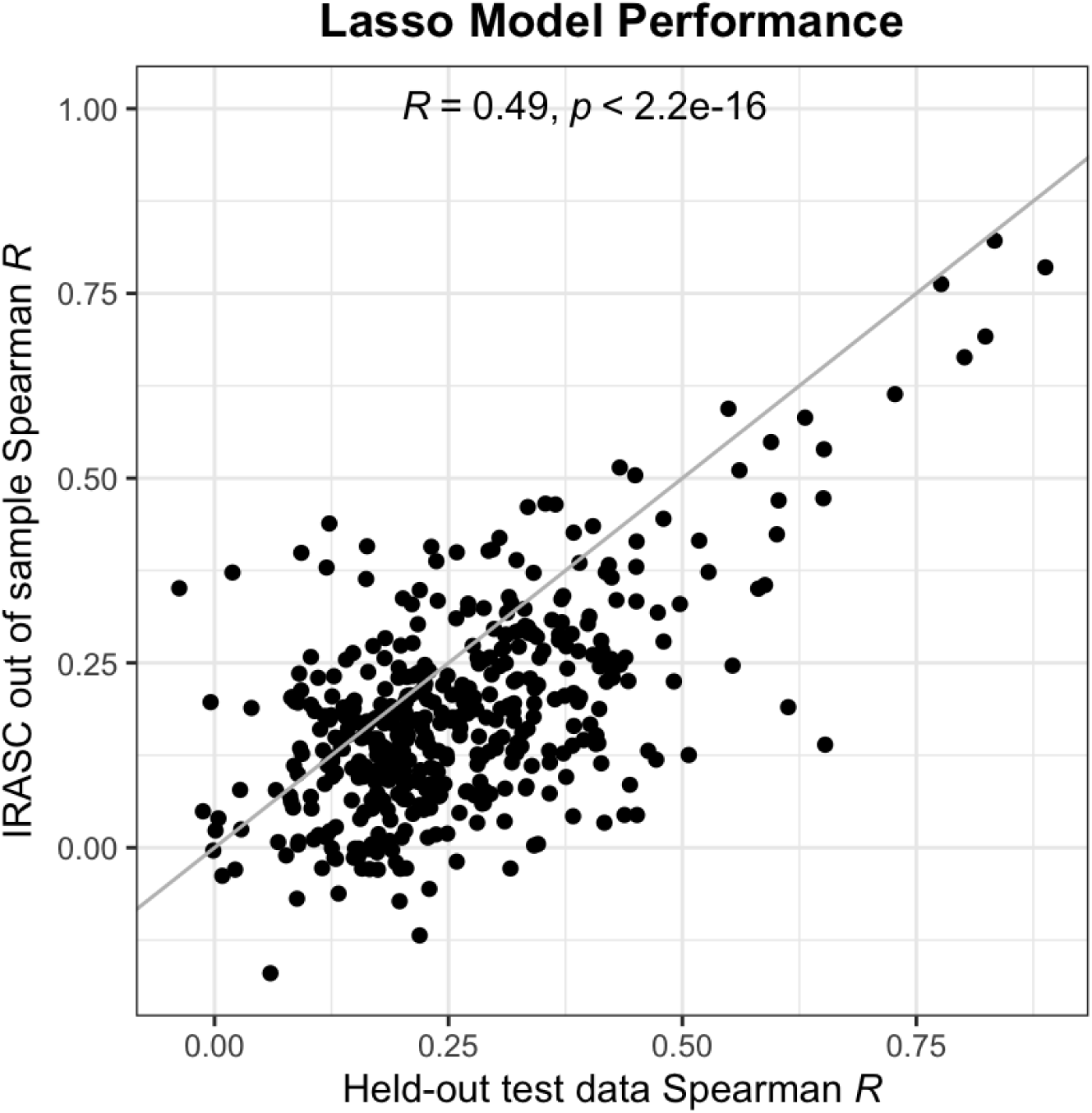
Concordance of the prediction performance across different testing cohorts. We performed predictions in two cohorts—held-out test dataset (METSIM, N=400) and out-of-sample dataset (IRASC, N∼200)—and compared them with the observed metabolite levels. Our models trained on the METSIM dataset (n = 5736) perform comparably between the held out test dataset and out-of sample-dataset.

**Figure S3:**
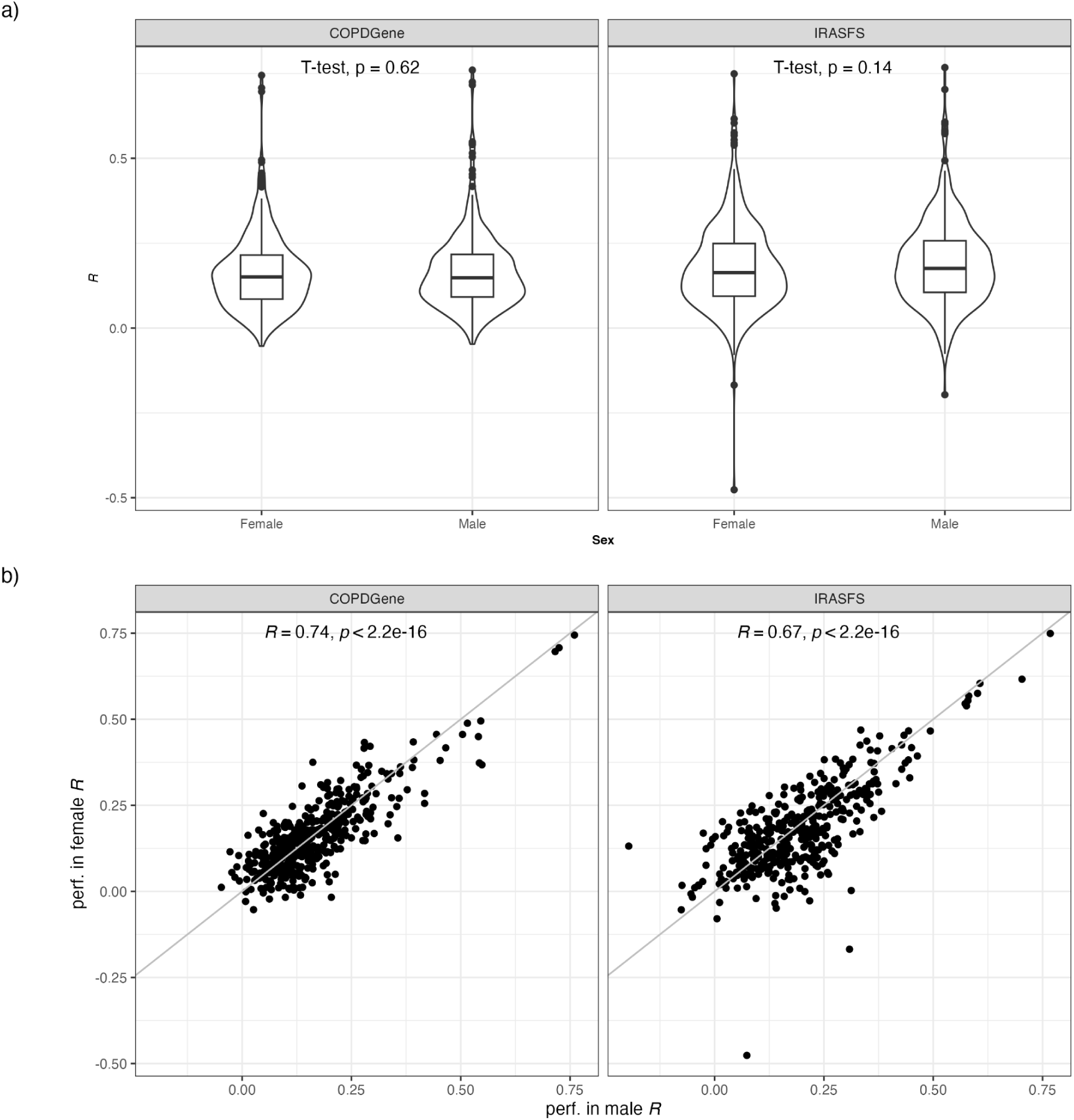
No obvious differences in performance between sexes in IRASFS and COPDGene test sets. Correlation between genetically predicted and observed metabolite levels in COPDGene individuals (left) with adjustment for covariates as described in Methods and IRASFS individuals (right). a) Distribution of Spearman correlation values across sex, b) Correlation of the prediction performance between genetically predicted and observed metabolite levels (i.e., Spearman correlation, R) in both sexes, indicating no obvious difference in prediction performance across sex in the COPDGene or IRASFS test data.

**Figure S4:**
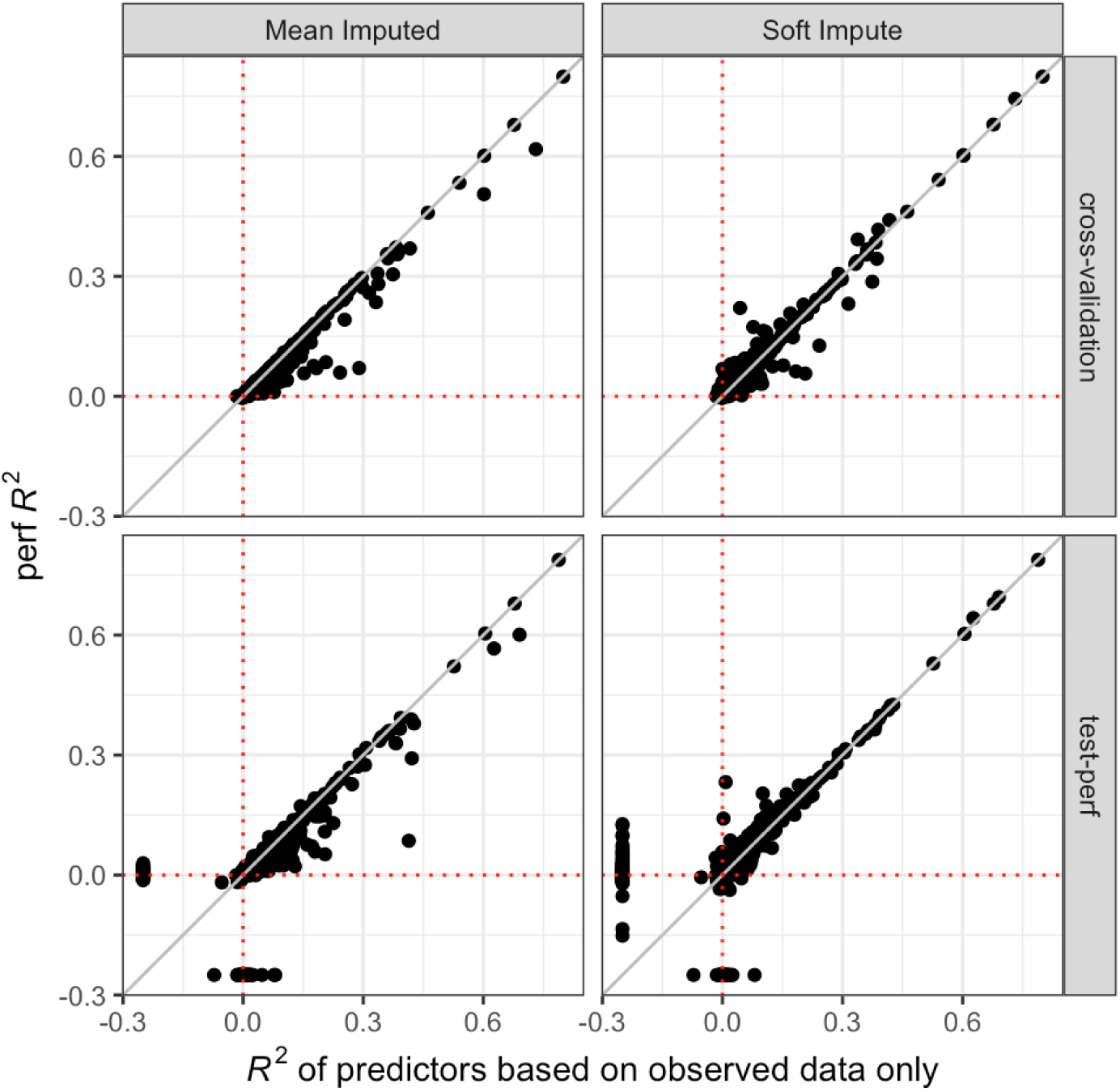
The softImpute method yields better predictors (larger R^2^) than mean imputation or no imputation (X-axis). Performance comparison of genetic predictors generated using different imputation methods (mean imputation and softImpute). We trained genetic predictors of metabolites on observed data (no missing values), mean imputed data, and soft imputed data and compared predictions from the models with the observed metabolite levels. We compared the performance of softImpute and mean imputed predictors against the predictors trained with no imputation. The top panels show performance in the cross validation dataset, and the bottom panels show performance in the held out dataset. The softImpute method performs better than the mean imputed and no imputation in 57% and 54% of the overlapping metabolites, respectively. The softImpute-based models have a median spearman correlation (R^2^) of 0.017 while the mean imputed data based models have an R^2^ of 0.013 in the held out test dataset.

**Figure S5:**
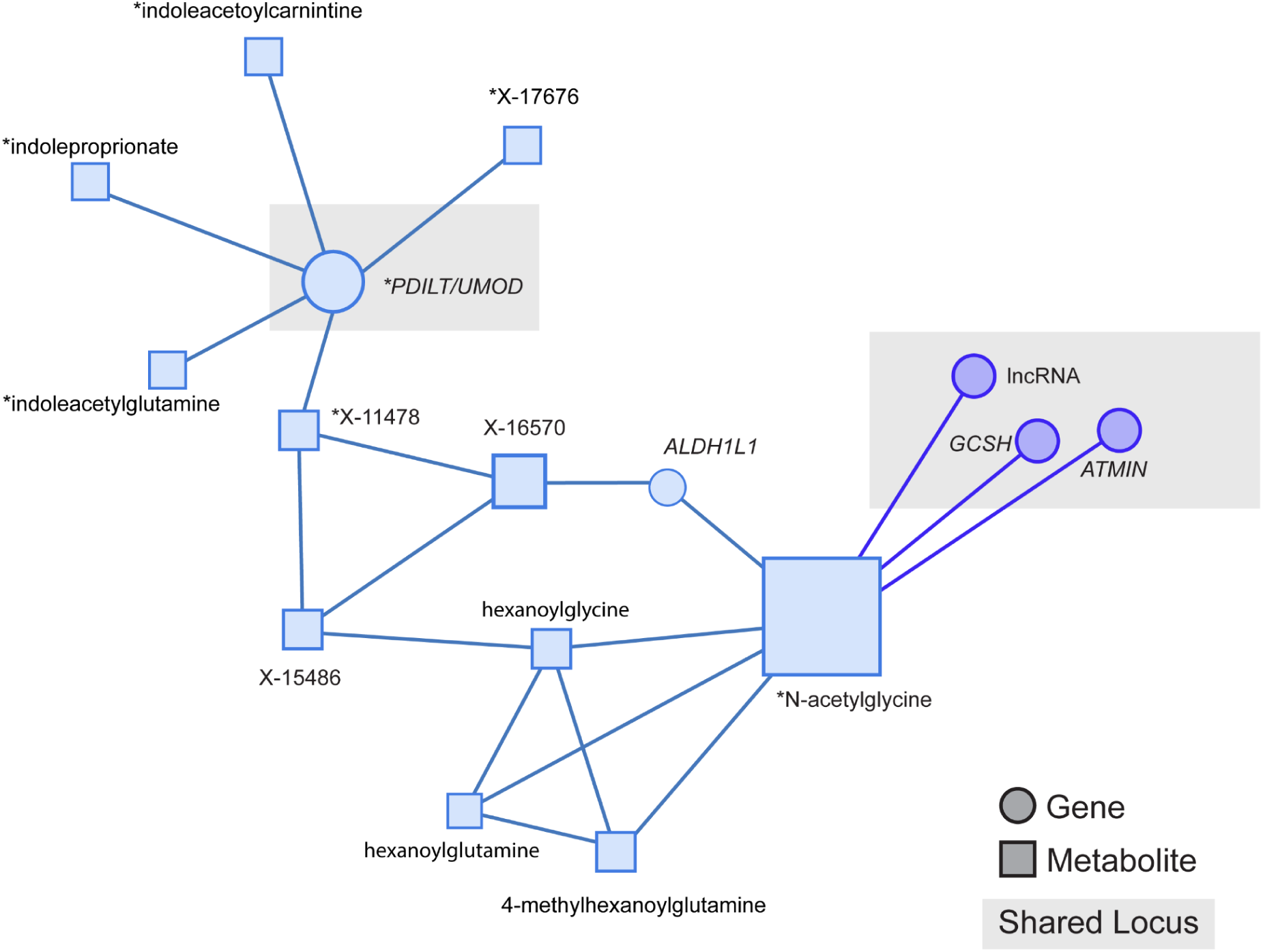
Portion of network for CKD showing connection between N-acetylglycine and PDILT locus. Inclusion criteria: i) all first-degree nodes for PDILT and UMOD, ii) all first-degree nodes for N-acetylglycine, and iii) nodes involved in shortest connection(s) between PDILT and N-acetylglycine. Genes are indicated as circles; metabolites as squares. * indicates individually significant genes and metabolites. Shape color indicates network membership, while shape size indicates the z-score.

### Supplemental Tables

**Table S1**: **Metabolite predictor training.** Metabolite model prediction performance, including lasso and ridge models, in both cross-validation and test datasets. This file also contains heritability estimates from GCTA software and metabolite annotations. https://docs.google.com/spreadsheets/d/15SvV9X1tMLmj1HKnQoipjXpmKBOWl2Pe/edit?usp=sharing&ouid=102425163912901090472&rtpof=true&sd=true

**Table S2**: MetaboXcan association results for CKD, which contains TWAS, MWAS, g-MWAS, pathways and network summaries.

